# Mechanism of negative *μ*-opioid receptor modulation by sodium ions

**DOI:** 10.1101/2023.06.15.545124

**Authors:** Neil J. Thomson, Ulrich Zachariae

**Affiliations:** Computational Biology, School of Life Sciences, University of Dundee, Dow Street, Dundee, DD1 5EH, United Kingdom; Biochemistry and Drug Discovery, School of Life Sciences, University of Dundee, Dow Street, Dundee, DD1 5EH, United Kingdom

## Abstract

The negative allosteric modulation of G-protein coupled receptors (GPCRs) by Na^+^ ions, known as the sodium effect, was first described in the 1970s for opioid receptors (ORs). Since then, it has been detected almost universally amongst class A GPCRs. High-resolution structures of class A GPCRs in the inactive state exhibit a Na^+^ ion bound to a conserved pocket near residue D2.50, whereas the active state structures of GPCRs are incompatible with Na^+^ binding. Correspondingly, Na^+^ ions diminish the affinity of receptor agonists, stabilize the receptors in the inactive state, and reduce basal signalling levels. Despite these observations, a detailed mechanistic explanation of how Na^+^ ions negatively modulate the receptor and inhibit activation has remained elusive. Here, we apply a mutual-information based analysis method to μs-timescale all-atom molecular dynamics simulations of the μ-OR to decipher conformational changes within the protein matrix and protein-internal water molecules that are directly coupled to the binding of Na^+^. Our results reveal that Na^+^ binding is tightly coupled to a water wire that links the Na^+^ binding site with the agonist binding pocket. Furthermore, Na^+^ binding leads to rearrangements in polar protein networks that propagate conformational changes both to the agonist and the intracellular G-protein binding sites via conserved micro-switch motifs. Our findings provide a mechanistic link between the presence of the ion and altered agonist binding affinity, receptor deactivation and the depression of basal signalling.

## Introduction

With over 800 members, G-protein coupled receptors (GPCRs) form the largest family of cell surface receptor proteins in humans. Embedded within the plasma membrane of the cell, the Golgi apparatus, and endosomes, these proteins act as signal transducers to enable transmembrane (TM) communication (Latorraca et al., 2017; Eichel and von Zastrow, 2018; Irannejad and Von Zastrow, 2014). At the cell membrane, extracellular agonist binding leads to receptor activation, involving conformational changes that expose effector protein binding sites on the receptors’ intracellular face. The effector proteins, including hetero-trimeric G-proteins and β-arrestins, initiate a range of intracellular signalling cascades that cause a wide variety of physiological changes in the cell. The diversity of the GPCR family means they are involved in an exceptionally broad spectrum of physiological functions, including vision, the detection of olfactory stimuli, neurotransmission, and immune responses (Martemyanov, 2014; Spehr and Munger, 2009; Venkatakrishnan et al., 2016). As such, GPCRs are the targets for two-thirds of all hormones and one-third of therapeutic drugs in the human body (Katritch et al., 2013; Hauser et al., 2018; Foster et al., 2019).

The class A (rhodopsin-like) GPCR subfamily includes the vast majority of GPCRs. They are characterised by a helical bundle structure comprising seven TM helices with connecting extracellular and intracellular loops, and a short membrane-parallel intracellular helix (H8) at the C-terminus (Rosenbaum et al., 2009; Katritch et al., 2013) (see Fig. 1). The ligand binding site for agonists, antagonists and inverse agonists is located in a deep water-filled pocket accessible from the exoplasm, while effector interactions are focused on a region encompassing the intracellular face of the helical bundle (Katritch et al., 2013; Fenalti et al., 2014; Koehl et al., 2018; Zhuang et al., 2022). Agonist-induced receptor activation is associated with a large-scale outward swing of the intracellular portion of TM6, opening access to this binding region for the effector G-protein (Koehl et al., 2018; Liu et al., 2019; Rose et al., 2014). Most GPCRs additionally exhibit a spontaneous level of basal signalling, independent of agonists. The inhibition of basal GPCR activity requires the binding of inverse agonists or sodium (Na^+^) to the receptors (Kobilka and Deupi, 2007; Katritch et al., 2014).

**Figure 1:**
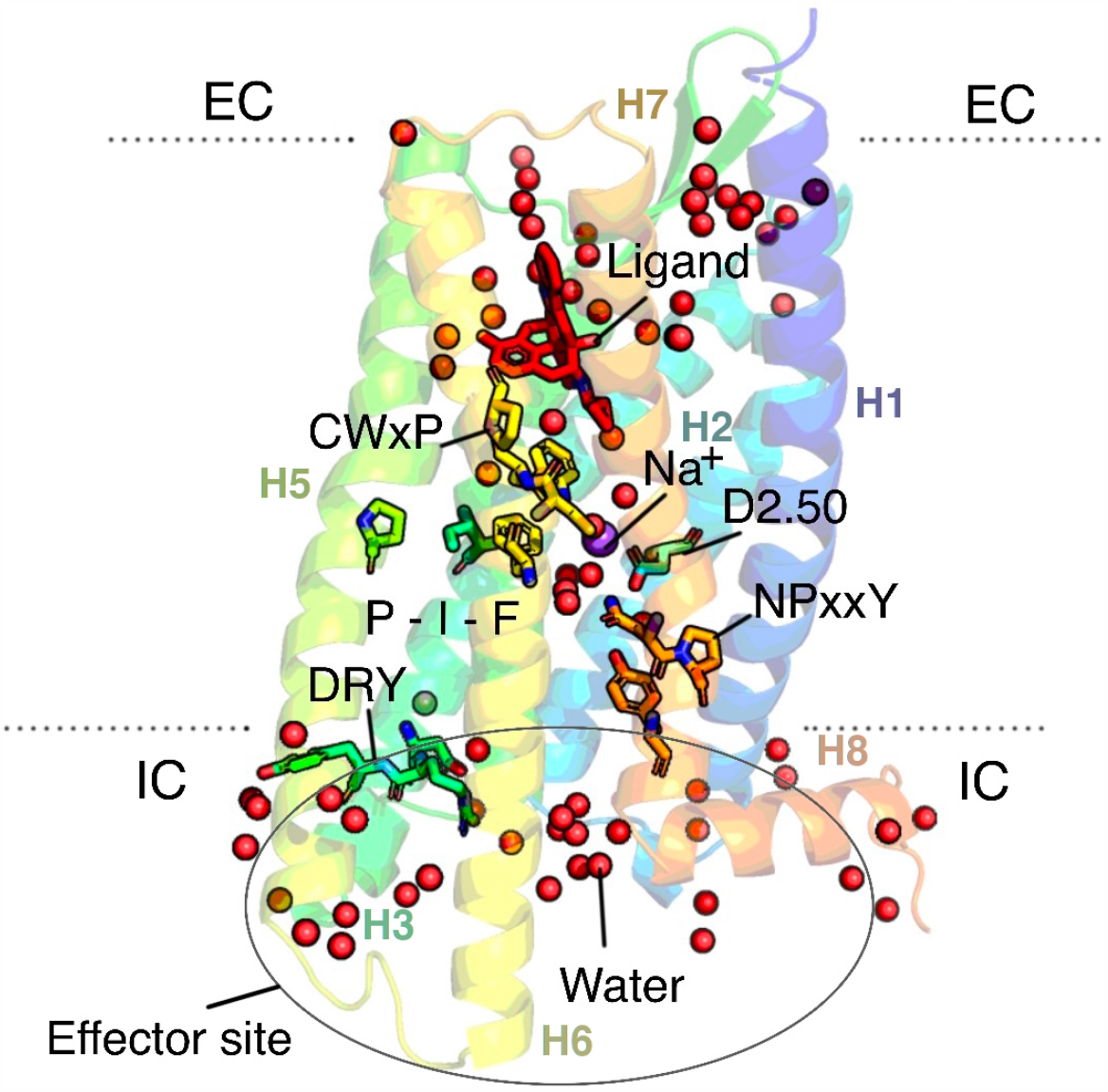
Key features of the μ-OR include: the seven transmembrane (TM) helical structure with an additional short, membrane-parallel helix at the intracellular (IC) domain; the orthosteric ligand binding site – accessible from the extracellular (EC) domain; the conserved CWxP, NPxxY, P-I-F and DRY motifs; the Na^+^-binding pocket and the primary Na^+^-coordination site – D2.50, and a diverse network of internal water molecules that mediate the internal polar network.

In the centre of the class A GPCR TM domain, multiple high-resolution crystal structures of antagonist-bound, inactive GPCRs reveal a Na^+^ ion bound to the highly conserved, ionizable aspartate residue D^2.50^ (Fenalti et al., 2014; Zarzycka et al., 2019) (receptor residues are numbered according to the Ballesteros-Weinstein scheme (Ballesteros and Weinstein, 1995; Isberg et al., 2016)). By contrast, structures of active receptors exhibit a collapsed Na^+^ binding site, unable to accommodate an ion. These structural findings link Na^+^ binding with receptor inactivation (Koehl et al., 2018; Katritch et al., 2014; Zarzycka et al., 2019). The negative allosteric modulation of GPCRs induced by Na^+^ was functionally evidenced 50 years ago, where it was shown that Na^+^ attenuated agonist binding in the class A opioid receptors, while antagonist binding was not significantly affected (Pert et al., 1973). This ‘Na^+^ effect’ has since been widely reproduced in other receptor types (Katritch et al., 2014; Friedman et al., 2020; Ferré et al., 2023). As well as diminishing agonist affinity, studies show that Na^+^ also depresses the level of agonist-independent basal activity in a wide range of GPCRs including the μ-opioid receptor (μ-OR) (Katritch et al., 2014; Quitterer et al., 1996; Selley et al., 2000). By contrast, mutations of D2.50 reduce the receptor’s affinity for Na^+^ and result in enhanced basal signalling (Katritch et al., 2014). The main Na^+^ binding site, its polar surroundings, and the protonation state of D2.50 have thus been suggested to play important roles in agonist-independent basal signalling of GPCRs (Zarzycka et al., 2019; Valentin-Hansen et al., 2015; Vickery et al., 2018). Furthermore, mutations of the Na^+^ binding site in the A_2A_ adenosine receptor were shown to abolish agonist-induced signalling and affect the conformations of key conserved amino-acid motifs implicated in GPCR activation, also termed ‘micro-switches’ (White et al., 2018). However to date, the mechanistic underpinnings of these wide-ranging functional effects of Na^+^ on GPCRs remain insufficiently understood.

The Na^+^ binding site in class A GPCRs at D2.50 is located within a highly conserved polar pocket, surrounded by residues L2.46, S3.39, N3.35, F6.44, W6.48, N7.45, S7.46, and N7.49 (Fenalti et al., 2014; White et al., 2018; Katritch et al., 2014; Kooistra et al., 2021). Notably, the pocket involves the highly conserved P5.50-I3.40-F6.44, CW6.48xP, and N7.49PxxY micro-switches, which undergo distinct conformational changes in the transition between inactive and active crystal structures (Venkatakrishnan et al., 2019; Fenalti et al., 2014; Koehl et al., 2018; Katritch et al., 2014; White et al., 2018) (Fig. 1). The ion is further embedded in a conserved network of water molecules that mediate a complex internal polar network with a distinct activation-state dependent fingerprint (Venkatakrishnan et al., 2016, 2019; Yuan et al., 2014; Katritch et al., 2013; Fenalti et al., 2014; Koehl et al., 2018). There is, however, a lack of information about how Na^+^ dynamically interacts with the internal distribution of water molecules and this polar network.

To shed light on these interactions, we investigated the effect of Na^+^ binding on the conformational states of the μ-OR including the water molecules that interconnect between conserved receptor motifs. We conducted repeated simulations of the μ-OR in two receptor conditions – the Na^+^-present and Na^+^-absent state – on a time-scale of ∼12 μs in total. All simulations were initiated from identical inactive μ-OR crystal structures including the co-crystallised antagonist (PDB ID: 4DKL). We did not make any changes to the nature (e.g. mutations) and protonation state of the Na^+^ binding site, in order to focus exclusively on the effect of the ion. To pinpoint conformational changes of protein residues and water molecules that are specifically linked to Na^+^ binding, we applied a mutual-information based analysis method termed State Specific Information, which detects conformational state transitions that are directly coupled to the presence of the ion (Vögele et al., 2022). Our analysis reveals an intricate functional interplay between the ion and receptor-internal water molecules, as well as distinct conformational changes within major protein micro-switches induced by the ion. By a two-part mechanism, these effects are propagated towards the agonist and main effector binding sites. Our findings explain both the reduction of agonist binding affinity and the reduction of signalling activity caused by Na^+^ in the μ-OR.

## Results and discussion

To characterise the interaction between protein-internal water molecules, receptor residue conformations, and Na^+^ binding, we first identified the location of water sites inside the protein matrix (see Methods). Subsequently, we used our recently developed mutual-information framework, State Specific Information (SSI, as implemented in the software PENSA (Vögele et al., 2022)) to identify regions of the receptor and its internal water network where conformational state changes are strongly coupled to the binding and unbinding of Na^+^. Because both the occupancy level of the water sites and the orientation of the water molecules therein may be relevant for receptor function, both features were included in our analysis.

*Na*^+^ *binding regulates a water chain between D2*.*50 and the agonist interaction site H6*.*52* Recent studies of the μ-OR have revealed great detail on the binding modes and affinities of a range of agonists (Vo et al., 2021; Qu et al., 2022; Zhuang et al., 2022; Mahinthichaichan et al., 2021). In specific, the potent opioid agonist fentanyl is known to engage in two distinct binding modes, discriminated by their interaction with the μ-OR residue H6.52. Mahinthichaichan *et al*. showed that fentanyl first engages with the μ-OR in a low affinity mode, associated with a positively charged H6.52, while deprotonation of H6.52 is required for fentanyl to move deeper into the agonist binding pocket and engage in a high affinity binding mode (Mahinthichaichan et al., 2021; Vo et al., 2021). To shed light on how Na^+^ binding might modulate agonist affinity, we first focused on internal water molecules and receptor residues in the vicinity of the orthosteric agonist binding pocket and the primary Na^+^ interaction site D2.50.

Our analysis identified four water sites bridging between D2.50 and H6.52 (termed W1–W4), in which water molecules form close hydrogen-bonding contacts and align between the two ionisable receptor residues (Fig. 2). In the Na^+^-bound state, the occupancy of W2 decreases by ∼40%, while all other water sites remain highly occupied in both states (Supplementary Table S2). We next probed the orientations of the bound water molecules by determining the polar coordinates of their dipole (see Methods and Supplementary Fig. S1) using PENSA (Vögele et al., 2022). Additionally, conformational changes of neighbouring protein residues were identified by monitoring backbone and sidechain dihedral angles. The analysis shows that the water molecules occupying sites W3 and W4 exhibit unique orientations in their binding cavities, distinctly coupled to the presence or absence of Na^+^ (Fig. 3). Amongst all residues and waters investigated, the orientation of the water molecule in site W4 is most strongly coupled to the binding of Na^+^ (in information-theoretic terms, W4 shares 0.65 bits of mutual information about whether Na^+^ is bound; for full SSI results see Supplementary Fig. S2 and Supplementary Table S2).

**Figure 2:**
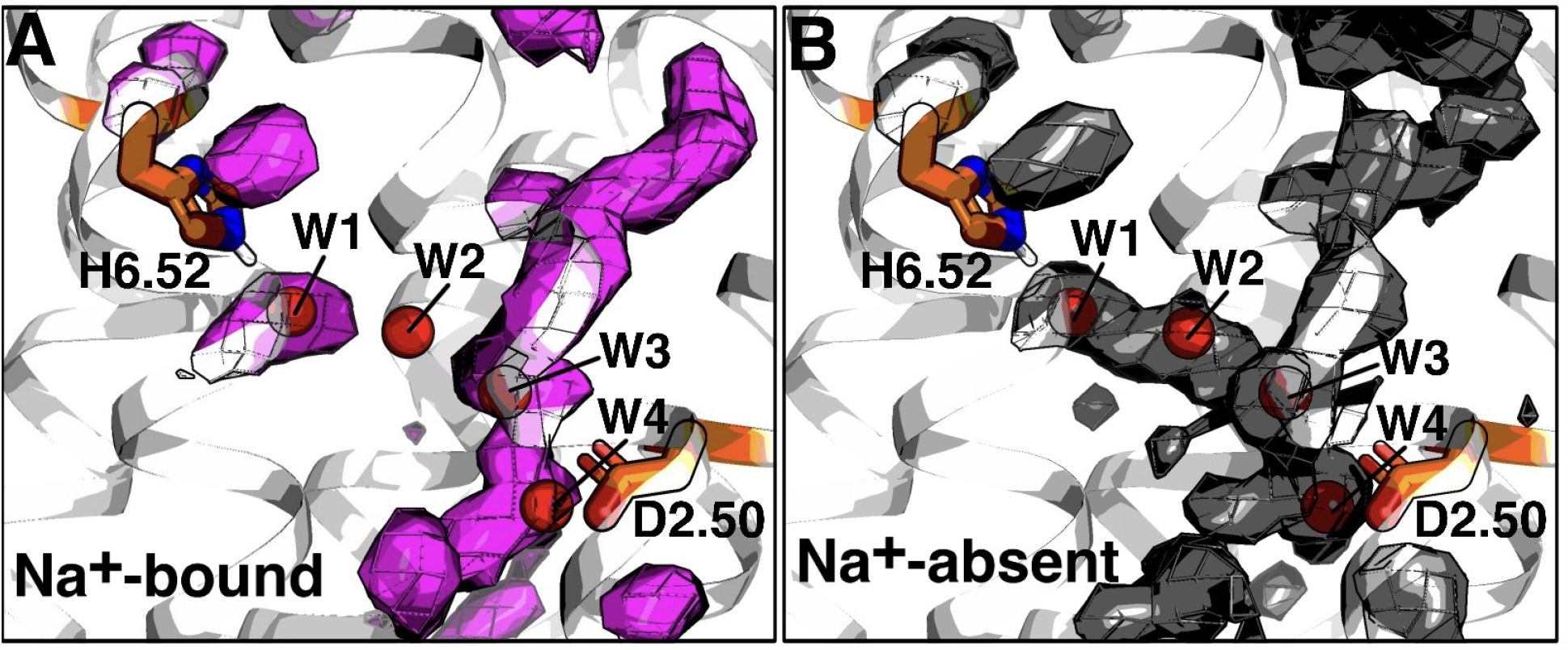
Water densities for Na^+^-bound simulations (A) and Na^+^-absent simulations (B) shown in mesh and surface representation at contour level 2. The water densities reveal that the hydration of a channel between H6.52 and D2.50 is modulated by Na^+^ binding. Red spheres represent the individual water cavities identified by PENSA, labelled W1, W2, W3 and W4.

**Figure 3:**
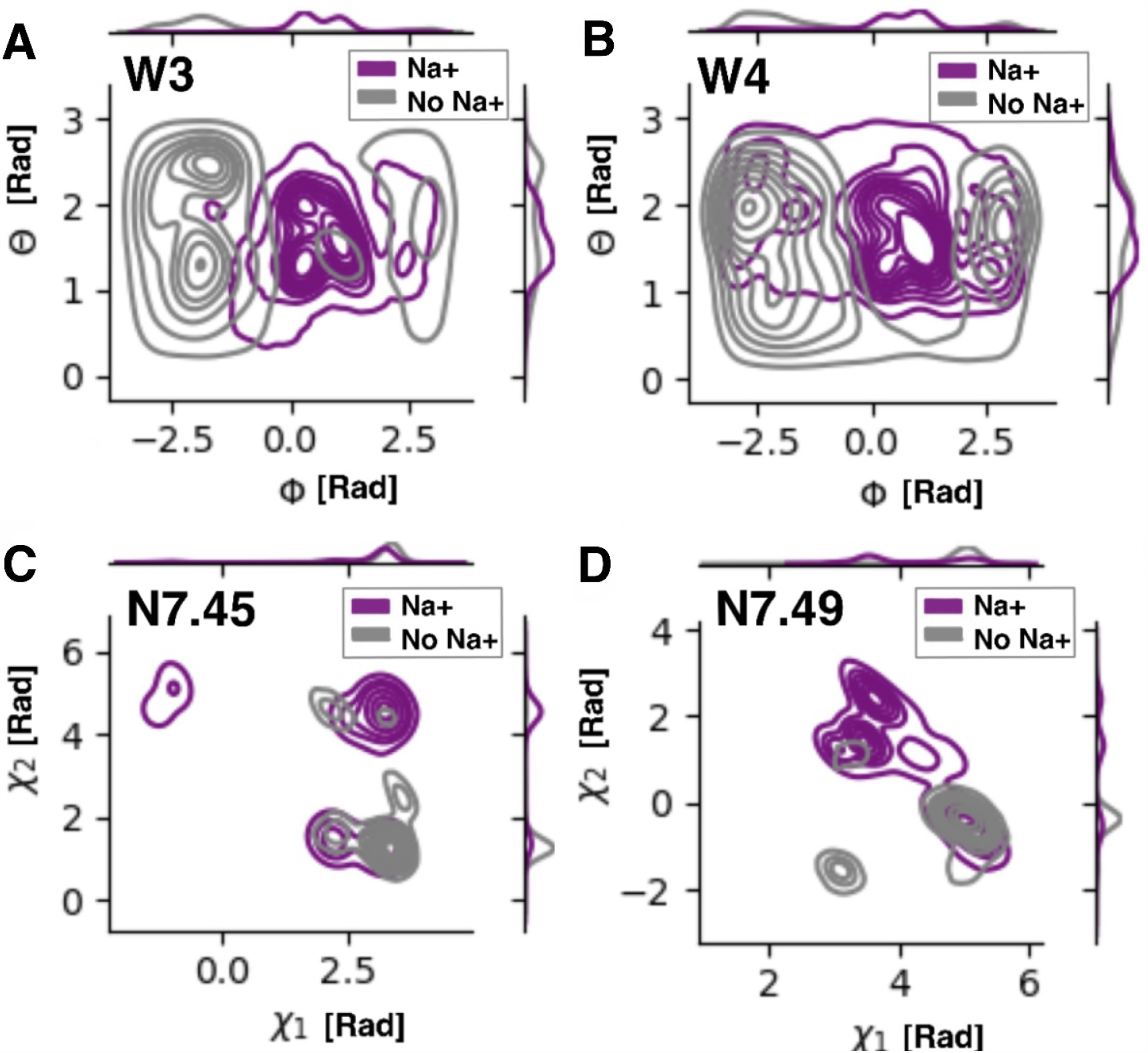
Analysis of internal water cavities reveals that Na^+^ modulates orientation of the water molecules in W3 (A) and W4 (B), and the sidechain rotamers of N7.45 (C) and N7.49 (D).

Taken together, the reorientation and occupancy changes within water sites W2–W4 serve to either establish or disrupt a water wire between the ionizable residues H6.52 and D2.50. In the absence of Na^+^, this water chain forms at a frequency of 0.39 ± 0.11. By contrast, Na^+^ binding entirely abolishes this water wire in our simulations (frequency of 0.0 ± 0.0). Accordingly, a continuous channel of water density is observed in the absence of Na^+^, whereas the water density displays a gap at water site W2 when the ion is present (see Fig. 2).

Furthermore, Na^+^ binding induces distinct sidechain orientations of the surrounding polar residues N7.45 and N7.49, which line the Na^+^ binding pocket and encompass water sites W3 and W4. Notably, N7.49 forms part of the N7.49PxxY micro-switch on TM7, a key GPCR motif implicated in receptor activation (White et al., 2018; Venkatakrishnan et al., 2019).

Deprotonation of H6.52, such that H6.52 adopts the neutral state, was shown to be a prerequisite for a high affinity fentanyl binding mode, while a second, lower-affinity binding mode occurred when H6.52 was protonated (Mahinthichaichan et al., 2021). Similarly, Vickery *et al*. showed that intracellular egress of the Na^+^ ion can occur upon receptor activation, and may be coupled to the protonation of D2.50 (Vickery et al., 2018). A spontaneously protonated D2.50 side chain may also underlie agonist-independent basal signalling (Vickery et al., 2018; Zarzycka et al., 2019) Concerted complementary protonation events at these two residues may thus be indicative of high affinity agonist-induced activation. Water wires, such as the four-molecule long chain reported here, can provide efficient avenues for proton transfer via a mechanism known as Grotthuss shuttling (Agmon, 1995). By preventing the formation of this possible proton transfer route, Na^+^ could maintain the positive state of H6.52 and prevent high affinity agonist binding. The water link might therefore serve as one explanation for the experimentally observed link between Na^+^ binding, reduced agonist affinity, and stabilisation of the μ-OR in the inactive state. However, the long-range effects of Na^+^ binding are not limited to protein-internal water molecules, as detailed below.

### Na^+^ binding directly impacts the interhelical packing between TM2, TM3, and TM6

Structural studies of the active μ-OR bound to a G-protein mimetic nanobody (Huang et al., 2015) and in complex with a G_i_-protein (Koehl et al., 2018) have highlighted that the active receptor state is characterised by a large outward swing of TM6, increasing its separation from TM3 and thereby exposing the G-protein binding site (Liu et al., 2019; Koehl et al., 2018). To shed further light on the connection between Na^+^ binding and diminished levels of both agonist-dependent and basal signalling, we therefore monitored packing changes between the receptor’s TM helices in the presence and absence of Na^+^. The distances between groups of Cα-atoms on each helix with similar coordinates along the membrane-normal axis were used to quantify inter-helical packing, as displayed in Fig. 4.

**Figure 4:**
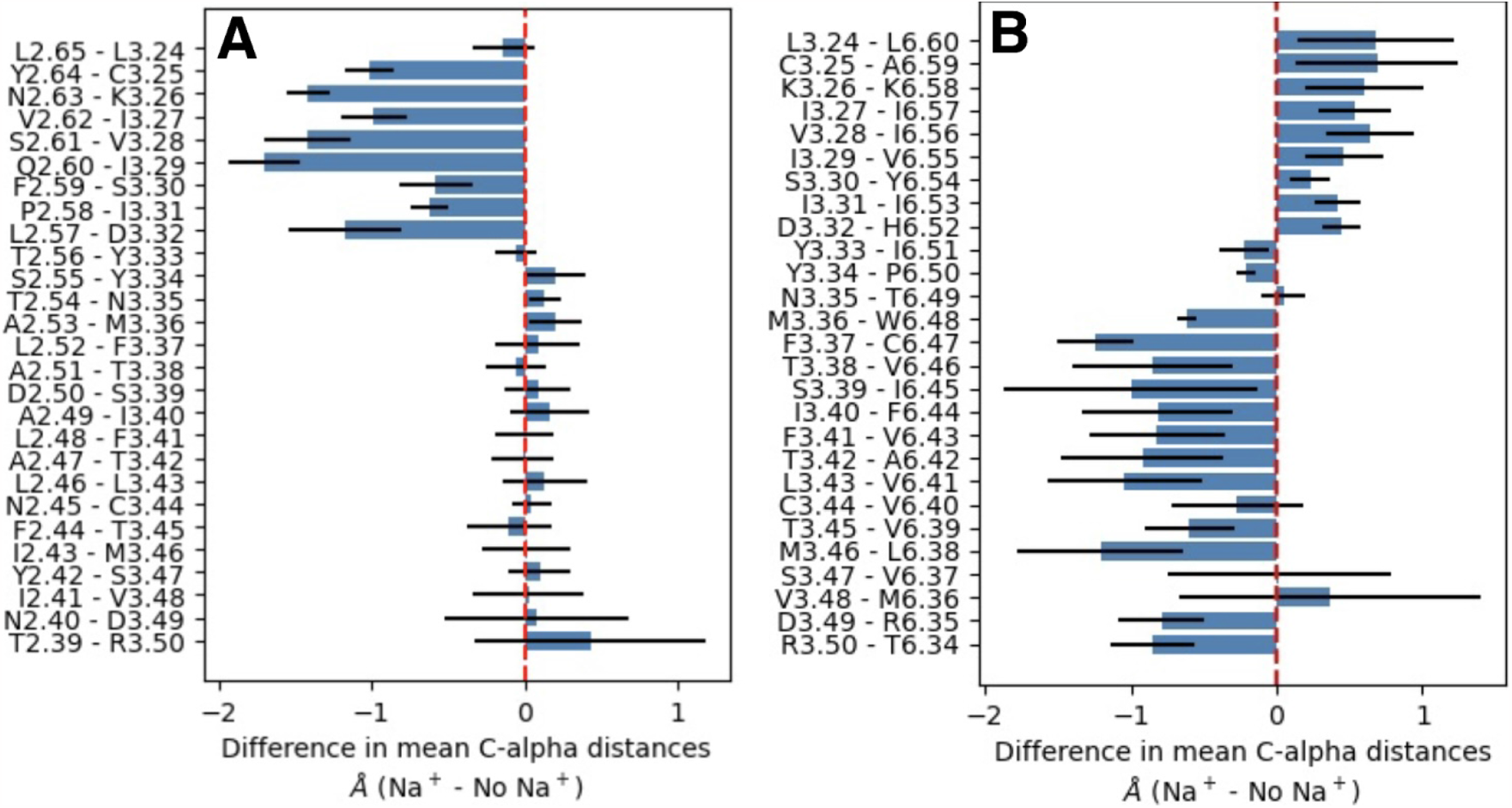
Inter-atomic distance analysis shows that Na^+^ modulates the inter-helical packing of TM3, represented by the difference in the mean Cα–Cα distances of residue pairs between each TM2 and TM3 (A), and TM3 and TM6 (B). Measurements below 0 indicate that Na^+^ increases the inter-helical packing, while measurements above 0 indicate that Na^+^ decreases the packing. Na^+^ increases the packing of TM3 against TM2 in the upper TM domain, while TM3 moves away from TM6 in this region. In contrast, TM3 packs closer against TM6 in the lower TM domain towards the G-protein binding site.

Our analysis shows that the packing of TM3 against TM2 and TM6 is significantly altered by the presence of the ion. In particular, Na^+^ induces a tighter packing between TM3 and TM6 from the Na^+^ pocket towards the intracellular domain, with the helices moving closer by ∼1 Å on average. In contrast, from the TM core towards the extracellular domain, the packing between TM3 and TM6 is loosened, instead increasing the compaction between TM3 and TM2, also by ∼1 Å on average (see Fig. 4).

Further investigation of the local configuration around the Na^+^ pocket shows that the compaction between TM3 and TM2 is caused by the ion forming tight interactions with N3.35, S3.39 and D2.50, which anchor TM3 to TM2. More specifically, the S3.39 and D2.50 sidechains are stabilised at a constant distance in the presence of Na^+^ (see Fig. 5). Similarly, the S3.39 sidechain hydroxyl group orients away from the ion and forms a strong hydrogen bond with the backbone oxygen atom of N3.35 at a distance of less than 2 Å (Fig. 4C-E). In contrast, absence of the ion destabilises the interactions between TM2 and TM3. The local reorganisation of N3.35, S3.39 and D2.50 on TM2 and TM3 is further propagated to TM6 by concerted motions of the conserved P-I-F and CWxP micro-switches, as detailed below.

**Figure 5:**
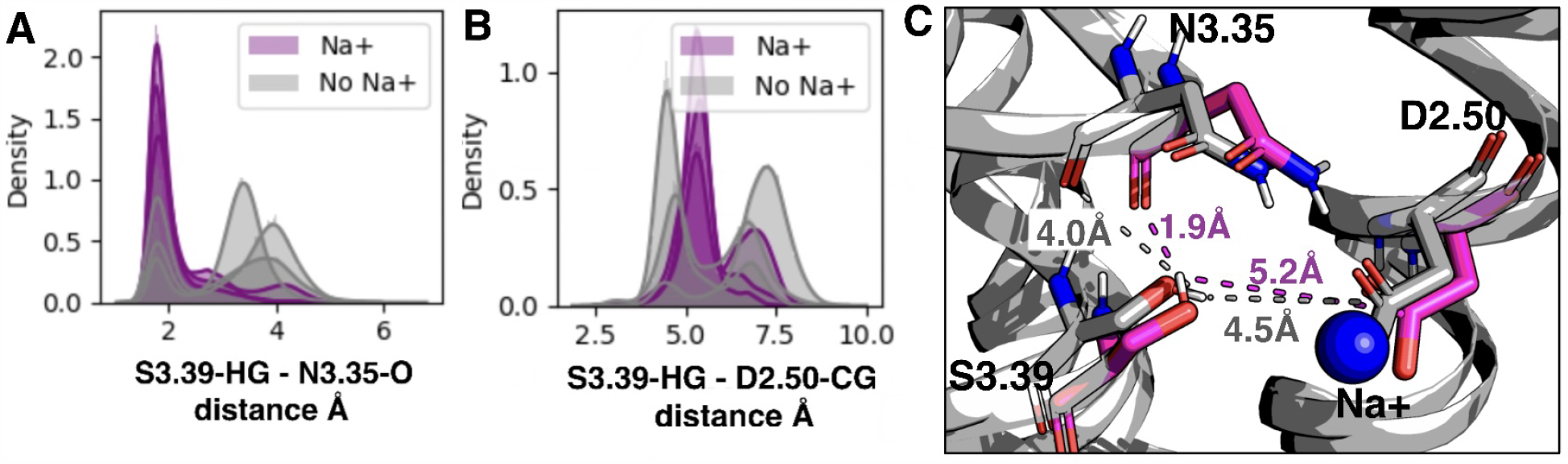
The increased packing of TM3 against TM2 in the presence of Na^+^ correlates with the formation of a strong hydrogen bond between the S3.39 hydroxyl and the N3.35 backbone (A), and the stabilisation of S3.39 with respect to D2.50 due to their shared Na^+^ interaction (B). The molecular conformation of the most likely interatomic distance distributions are shown in the panel C.

### Na^+^ binding leads to conformational changes at the effector binding site linked to reduced activity

As previously noted, the packing between TM3 and TM6 is a crucial determinant of receptor activity. In this context, the DRY motif on the intracellular face of TM3 plays a particularly important role in the outward movement of TM6 upon activation and in G-protein binding. Ionic locks formed between the charged residues D3.49 and R3.50 of the DRY motif and between R3.50 and E6.30 on TM6 are characteristic of inactive class A receptors, as they maintain a close contact between TM3 and TM6 and keep the G-protein binding cavity in a closed state (Katritch et al., 2013; Ballesteros et al., 2001; Dror et al., 2009; Liu et al., 2019). Accordingly, mutations of D3.49 and E6.30 have been shown to significantly increase constitutive activity in a variety of receptor types (Hulme, 2013; Kobilka and Deupi, 2007; Alewijnse et al., 2000). By contrast, when R3.50 is freed from the ionic locks, the R3.50 sidechain serves as a primary coordination site for the signal-transducing G-protein (Huang et al., 2015; Koehl et al., 2018; Liu et al., 2019).

Even though the R3.50—E6.30 ionic lock is absent in the ORs (due to a leucine residue in place of the glutamate), our analysis shows that a similar R3.50—L6.30 ‘polar lock’ is formed in the same region of the μ-OR between the sidechain guanidinium group of R3.50 and the backbone oxygen atom of L6.30. We find that the formation of this polar lock correlates with the presence of Na^+^ via the tightening of the broader TM3-TM6 interaction described above. The locked state is favoured in the presence of the ion, while the unlocked state is promoted by its absence (see Fig. 6). Na^+^ binding thus both inhibits the outward swing of TM6 associated with receptor activation (Katritch et al., 2013) and prevents the R3.50 sidechain from rotating towards TM7, the rotameric state associated with the docking of G-proteins (Huang et al., 2015; Koehl et al., 2018; Liu et al., 2019). This result can serve as a further explanation for the observed negative allosteric modulation of the μ-OR by Na^+^ ions.

**Figure 6:**
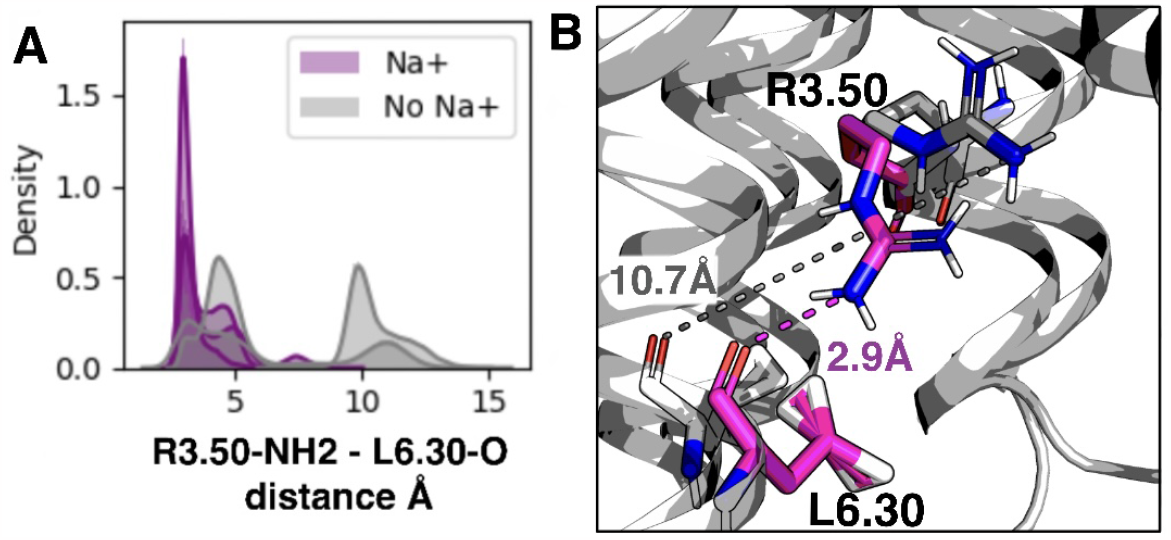
The increased intracellular packing of TM3 against TM6 in the presence of Na^+^ correlates with the formation of a strong hydrogen bond between the R3.50 sidechain amine and the L6.30 backbone (A, B). The molecular conformation of the most likely interatomic distance distributions are shown in the panel B.

**Figure 7:**
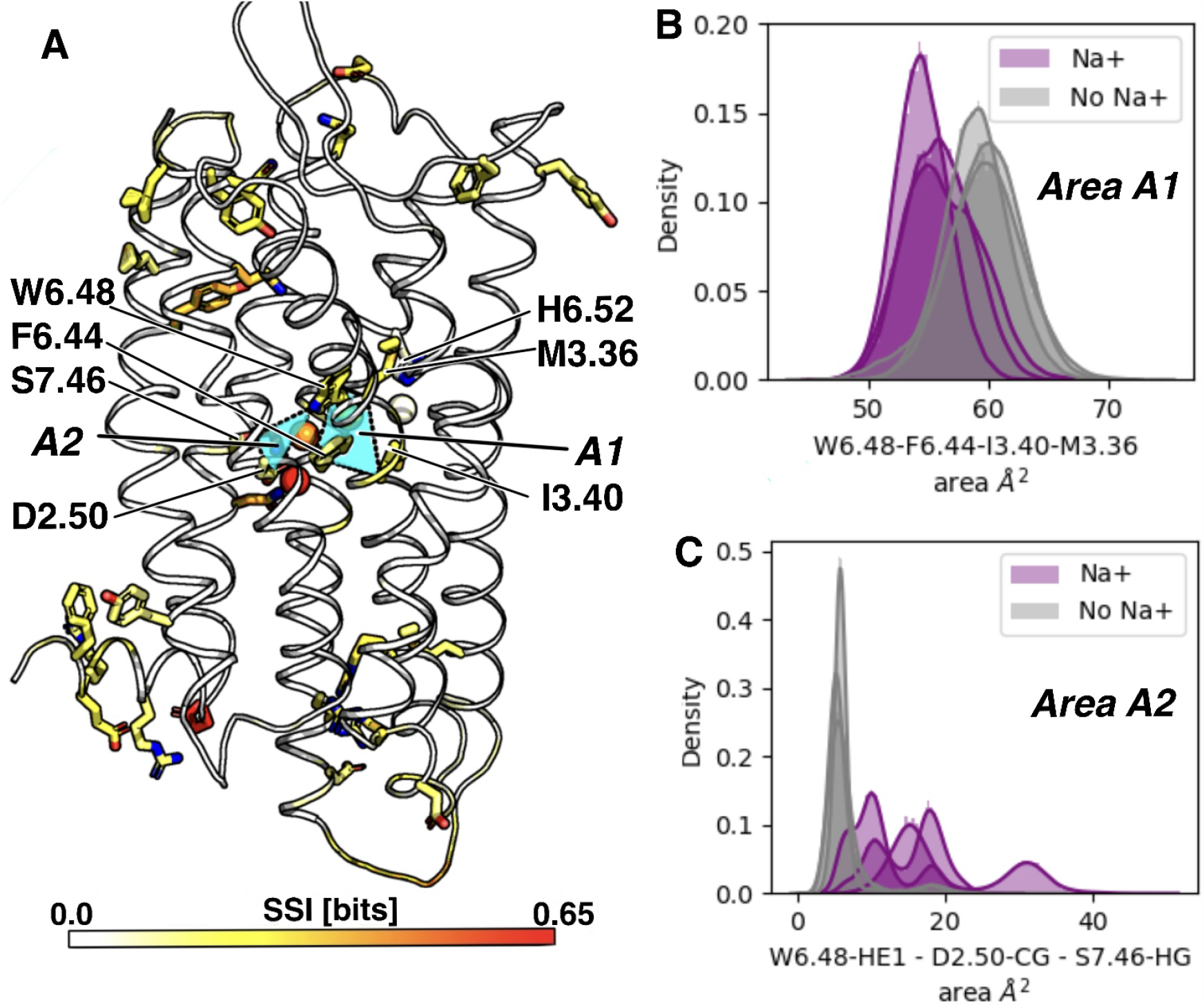
(A) State-Specific Information (SSI) for all residue backbone (cartoon representation) and sidechain (stick representation) torsion angles, and for intracavity water sites (orientation and occupancy, sphere representation). The SSI quantifies the magnitude of coupling between Na^+^ binding events and changes to the conformational sampling of each feature. SSI values are projected onto the μ-opioid structure in a colour code ranging from white (0 bits) to red (0.65 bits) through yellow and orange. To highlight the most closely coupled sites, only residues with an SSI value greater than 0.1 are shown in stick representation; H6.52, D2.50 and S7.46 are further displayed for reference. Na^+^ binding increases the packing between the hydrophobes W6.48, F6.44, I3.40 and M3.36, reflected by a reduction in the area between these residues’ Cα atoms (B). In contrast, in the absence of Na^+^, the altered polar network in the ion binding pocket results in W6.48 swinging into a defined conformation towards S7.46 and D2.50, increasing the packing between these three residues and altering the shape of the agonist binding site (D).

**Figure 8:**
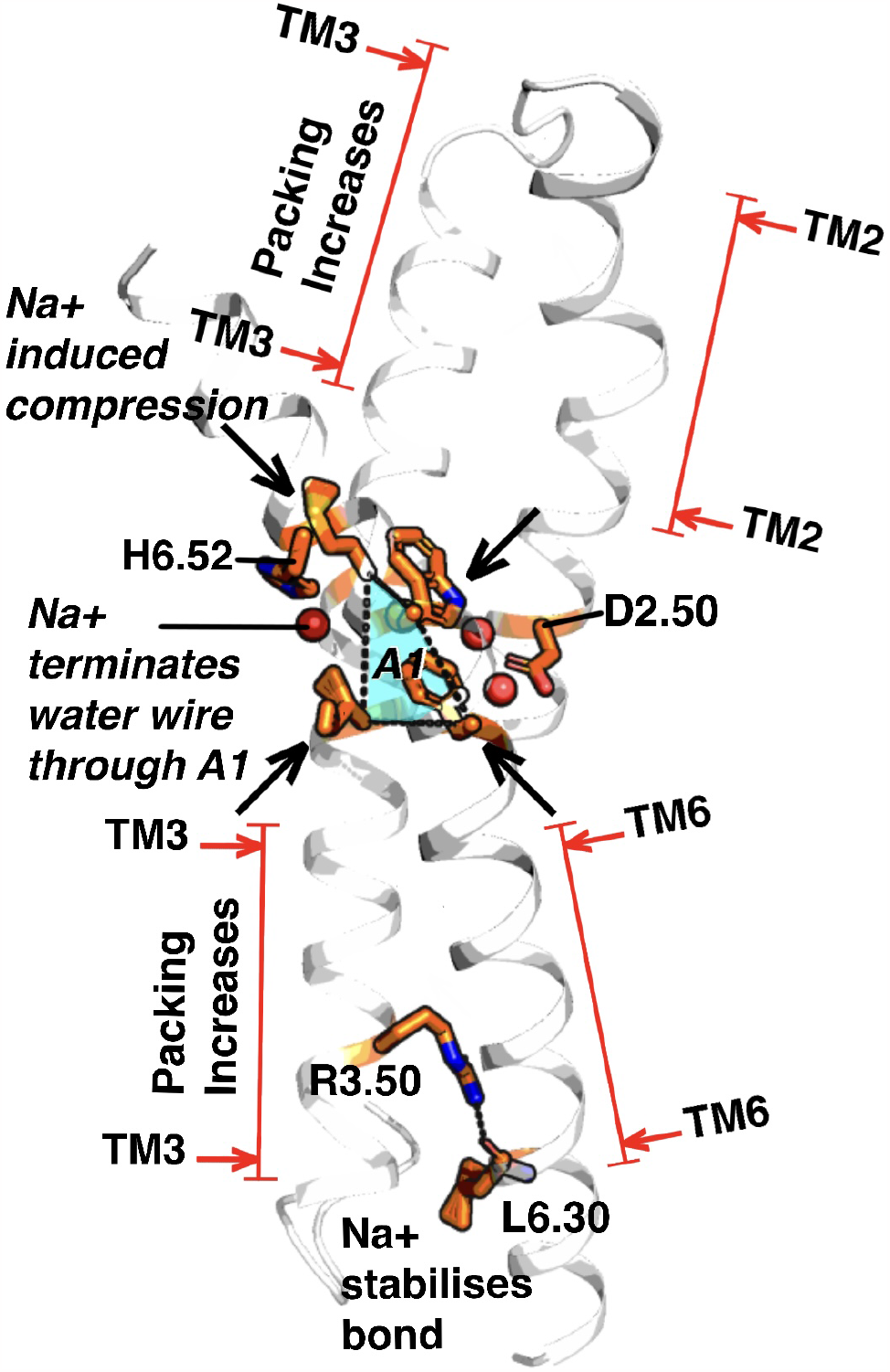
Na^+^ binding increases the packing between the hydrophobic residues W6.48, F6.44, I3.40 and M3.36, resulting in the termination of a water connection between H6.52 and D2.50, which traverses through the area enclosed by the hydrophobes. The hydrophobic packing increase propagates down TM3 and TM6 towards the intracellular domain, where the two helices are stabilised at close proximity by the formation of a polar bond between R3.50 (DRY motif) and the L6.30 backbone. Similarly, the upper portion of TM3 forms a tighter packing against TM2 in the extracellular region, affecting further the conformation of the ligand binding site.

### Concerted conformational changes in the P-I-F and CWxP motifs propagate the local Na^+^ effects across the receptor

Since Na^+^ binding triggers a substantial long-range repacking of TM6, TM3, and TM2, we were next interested in the coupling mechanism between the local Na^+^ effects and these distal conformational changes. Our SSI analysis shows that the sidechain conformational states of W6.48 (CWxP motif), F6.44 (P-I-F motif), I3.40 (P-I-F motif), and M3.36 are all affected by Na^+^ binding (for full SSI results, see Supplementary Fig. S2). These residues, mostly large hydrophobes, are situated one helix turn apart on helix TM6, and opposite each other between TM6 and TM3, respectively. Both motifs have previously been identified as major functionally relevant protein micro-switches, in which packing changes between W6.48 and F6.44, and between W6.48 and I3.40, are indicative of receptor activation in a variety of receptor types (Martí-Solano et al., 2014; Smith, 2021; Rasmussen et al., 2011; Zhou et al., 2019). We therefore hypothesised that the Na^+^ ion might affect the hydrophobic interactions in this region and used the area of a quadrilateral formed between the four sidechains to quantify their hydrophobic packing level.

We find that the area between the Cα atoms of W6.48, F6.44, I3.40 and M3.36 decreases by ∼4 Å^2^ upon Na^+^ binding, from 59.4±0.2 Å^2^ to 55.5±0.5 Å^2^. In addition, these four residues form a perimeter around the water connection between H6.52 and D2.50 addressed above, with water site W2 locating almost exactly to the geometric centre of the residues’ Cα atoms. The increased packing of residues W6.48, F6.44, I3.40 and M3.36 is directly correlated with the reduction of water occupancy in water site W2, since this compaction diminishes the volume that is available for a water molecule. Furthermore, the tightened interactions made by W6.48, F6.44, I3.40, and M3.36, are concerted with the increased packing of TM6 and TM3 towards their intracellular side, implicating this compaction in the global interaction between TM6 and TM3.

By contrast, a rearrangement and strengthening of interactions in a surrounding polar network occurs in the absence of Na^+^. Without Na^+^, W6.48 in the orthosteric ligand binding pocket orients towards S7.46 and D2.50, forming a stable hydrogen bonding interaction. This is reflected by a reduction of over 50% in the area between W6.48—HE1, D2.50—CG and S7.46—HG from 15.5±2.8 Å^2^ to 6.4±0.2 Å^2^. The tightened polar interaction network disrupts the hydrophobic packing of the CWxP and P-I-F motifs by reorienting the W6.48 sidechain away from I3.40, M3.36 and F6.44. The thereby increased polar interaction also correlates with the formation of the water connection between H6.52 and D2.50.

Residue W6.48, which forms the floor of the agonist binding pocket, thus switches its sidechain between a conformation with strengthened hydrophobic interactions in the presence of Na^+^, and a conformation which enhances the formation of a polar network in its absence. Whereas Na^+^ binding to D2.50 absorbs many of the polar functional groups required to establish this polar network, they are released when the ion binding pocket is vacant. In this way, Na^+^ alters the shape of the ligand binding pocket of the μ-OR. Agonists such as fentanyl show direct contacts with residue W6.48 in structures of the active state μ-OR (Zhuang et al., 2022). It is therefore likely that a conformational shift of the W6.48 sidechain has a further impact on agonist affinity.

### Relevance for GPCR basal signalling

In addition to stabilising the receptor inactive state, Na^+^ has been experimentally shown to reduce the basal signalling level of the μ-OR (Selley et al., 2000). As it is likely that basally active receptors display a significant proportion of the R3.50 sidechain population in the unlocked state, competent to bind G-proteins, irrespective of agonist binding, our findings above suggest a mechanism which provides a ligand-independent level of control of this sidechain by Na^+^, maintaining it in the locked conformation.

Moreover, they explain mutagenesis studies in which alanine substitutions at S3.39 and N3.35 increase constitutive activity (Hulme, 2013), as these mutations would disrupt the coupling mechanism proximal to the Na^+^ binding site. Mutations of S3.39 and N3.35 also confer constitutive activity to the otherwise basally quiescent neurokinin receptor NK1 (Valentin-Hansen et al., 2015). In the wild-type NK1 receptor, the otherwise highly conserved Na^+^ binding residue D2.50 is replaced by E2.50, and the longer glutamate sidechain occupies the binding space of the Na^+^ ion (Zarzycka et al., 2019; Valentin-Hansen et al., 2015; Schöppe et al., 2019).

The basally quiescent wild-type NK1 receptor could thus be considered as a GPCR with a permanently occupied Na^+^ binding pocket, abolishing basal activity via the connectors S3.39 and N3.35. Taken together, our results provide a mechanistic explanation for the widely observed negative Na^+^ effects on GPCR agonist-induced and basal activity, and the role of a range of constitutively activating mutations, locating to the coupling and end points of this mechanism.

## Conclusion

The inhibitory Na^+^ effect on GPCR activation was first described for ORs in 1973 (Pert et al., 1973). Although the Na^+^ effect has been detected in many other receptor types since, the mechanistic underpinnings of the typical negative modulation of GPCRs by Na^+^ have remained insufficiently understood. Our findings, obtained through a mutual-information based analysis of μs-timescale atomistic simulations of the μ-OR, reveal how the local effects of Na^+^ binding are propagated to both the agonist and the effector G-protein binding sites, by a mechanism involving both functionally relevant protein micro-switches and protein-internal water molecules. The results therefore highlight the importance of a coupling mechanism between bound ions, water molecules, and the protein matrix for protein function.

The Na^+^ effect is spread across the entire TM section of the μ-OR by a two-part mechanism in which Na^+^ binding initially anchors S3.39 and N3.35 to D2.50, stabilising the conformations of the two TM3 residues close to TM2. The tightening prevents the agonist-binding residue W6.48 from swinging away from TM3 and towards TM7 and thereby leads to an increased hydrophobic packing between M3.36, I3.40, F6.44, and W6.48 of the CWxP and P-I-F motifs. This repacking is propagated to the intracellular side of TM3 and TM6 including the main effector binding region at R3.50 and the DRY motif. In addition, it terminates a water connection between the ionizable agonist binding residue H6.52 and the ionizable Na^+^ binding residue D2.50, where equal and opposite protonation events are potentially characteristic of high affinity agonist-induced activation (Mahinthichaichan et al., 2021; Vickery et al., 2018).

All of these rearrangements are ultimately due to the changed hydrogen bonding pattern at and around the Na^+^ binding site D2.50, while the key movements propagating the Na^+^ effect to the ligand and effector binding sites are centred on TM6. The mechanism we observe can thus serve to explain the alterations of both the agonist affinity and the signalling activity of the receptor upon Na^+^ binding.

These results suggest that replication of the Na^+^ effect with, for example, a bitopic ligand may reduce the binding affinity of fentanyl. Correspondingly, Faouzi *et al*. recently developed a bitopic fentanyl derivative which targets both the Na^+^ binding site and the conventional orthosteric fentanyl binding site (Faouzi et al., 2023). This bitopic fentanyl derivative binds in the low affinity mode identified by Mahinthichaichan *et al*., and was shown to maintain a high G_i_ potency with reduced arrestin recruitment (Mahinthichaichan et al., 2021; Faouzi et al., 2023). Taken this together, this implicates distinct binding modes and their modulation by receptor protoination events in biased signalling, however further work would be needed to verify this.

## STAR Methods

### Key Resources Table

| REAGENT | RESOURCE SOURCE | IDENTIFIER |
| --- | --- | --- |
| Software and algorithms |  |  |
| GROMACS | Abraham et al. (2015) | https://www.gromacs.org |
| PENSA | Vögele et al. (2022) | https://github.com/drorlab/pensa |
| MDANALYSIS | Gowers et al. (2016) | https://www.mdanalysis.org |
| ACPYPE | Sousa Da Silva and Vranken (2012) | https://www.bio2byte.be/acpype |

#### Contact for Reagent and Resource Sharing

Further information and requests for resources and reagents should be directed to and will be fulfilled by the lead contact, Ulrich Zachariae.

### Method Details

#### Molecular dynamics simulations

All molecular dynamics simulations were performed using the software GROMACS, version 5.1 (Abraham et al., 2015; Lindahl et al., 2020). The simulations were based on the crystal structure of the inactive μ-OR (PDB entry 4DKL) (Manglik et al., 2016). The refined structure was selected from GPCRdb (Kooistra et al., 2021) to ensure any mutations in the crystallographic protein construct were reversed to the canonical sequence. The receptor was truncated and capped with acetyl and N-methyl moieties and the N and C terminals, respectively. All crystal waters in the TM domain were left within the receptor, and the receptor was embedded in a homogenous 1-palmitoyl-2-oleoyl-glycero-3-phosphocholine (POPC) membrane bilayer, and solvated and neutralised in 150 mmol/L NaCl solution. For POPC, we used the SLipid model (Jämbeck and Lyubartsev, 2012), for water, the TIP3P model (Jorgensen et al., 1983), and for the protein, the amber99sb-ildn forcefield was selected (Lindorff-Larsen et al., 2010). Virtual sites were employed to enable an integration timestep of 4 fs (Feenstra et al., 1999). The ion parameters were adopted from the work of Joung and Cheatham (Joung and Cheatham, 2008). Three independent simulations were performed for each condition.

The bound antagonist was parameterised using ACPYPE (Sousa Da Silva and Vranken, 2012) and its charge state verified. The size of the membrane and simulation box were selected such that the receptor’s short-range Coulombic interactions across the periodic boundary were outside the range of self-interaction. The simulations were initially minimised with the steepest descent method (Meza, 2010). Subsequent equilibration in the NVT and NPT thermodynamic ensembles was performed for 10 ns each. The pressure was set to 1 bar and maintained constant with Parrinello-Rahman semi-isotropic pressure coupling (Parrinello and Rahman, 1981) using a time constant of 5 ps. Similarly, the temperature was maintained at a physiological level of 310 K with the Nosé-Hoover thermostat (Evans and Holian, 1985). The receptor-ligand complex, membrane, and solvent were all independently coupled to the temperature bath. A linear re-scaling was applied between the receptor-ligand-membrane subsystem and the solvent to remove any motion of the solvent relative to the bilayer. Long-range electrostatic interactions were modelled using the particle-mesh Ewald method (Darden et al., 1993) with a cut-off of 12 Å. The LINCS algorithm (Hess et al., 1997) was used to constrain bond lengths.

#### Definition of water sites and rotamer states

The water sites were determined from the simulations’ water density grid using PENSA (Vögele et al., 2022). Water sites are defined as spheres of radius 3.5Å, centred on the water density probability maxima, as described in (Vögele et al., 2022). A timeseries distribution is obtained for each water site by first identifying for each frame in the simulation’s trajectory whether the sphere (water site) is unoccupied or occupied. If occupied, the orientation of the water molecule is obtained by converting the water’s dipole moment into the angular components of spherical coordinates. The water site timeseries is then a combination of the spherical coordinates and a value outside the phase space of the spherical coordinates to represent an unoccupied state. For those frames where more than one water molecule occupies a water site, the dipole moment of the water molecule most frequently occupying the site is used to represent the orientation for that frame.

The water site centres can be in close enough proximity that the volumes used to calculate the water site occupancy statistics overlap. Measurements of a water site’s occupancy may therefore be influenced by the neighbouring sites. To account for this limitation in those cases, we also visually analysed all trajectory’s water density grids, confirming the reported water occupancy changes were present (see Fig. 2).

Rotamer states for receptor backbone sites comprise of the ϕ and ψ torsion angles, combined into a multivariate state, as described in (Vögele et al., 2022). Similarly, all sidechain torsion angles are combined to represent the sidechain conformation. The resulting State-Specific Information values therefore refer to the entire (geometric) conformational space of each residue.

#### Water wire analysis

To measure the water wire formation frequency, the MDAnalysis water bridge analysis tool was used (Gowers et al., 2016; Michaud-Agrawal et al., 2011). This analysis recognises neighbouring waters as a wire if they align such that the oxygens on one water are within hydrogen bonding distance of the hydrogens on the neighbouring water.

## Acknowledgments

NJT and UZ were supported by the UKRI Biotechnology and Biological Sciences Research Council (BBSRC) [grant number BB/M010996/1]. We thank David Gray and Andrei Pisliakov for helpful comments on this work.

## Author Contributions

NJT and UZ conceived and designed the research, NJT performed simulations, wrote analysis code and analysed the data, NJT and UZ wrote and edited the manuscript, UZ supervised the work.

