## Supplemental information for "Mechanism of negative *μ*-opioid receptor modulation by sodium ions"

**Table S 1.** Frequencies for which each water site is occupied by a water molecule under both simulation conditions – $Na^+$-bound and $Na^+$-absent. Since hydrogen bonds may be maintained while water molecules move around within a cavity, cavities are considered occupied if a water molecule exists within a 3 Å radius of the cavity centre.

| | W1 | W2 | W3 | W4 |
|---|---|---|---|---|
| $Na^+$-bound | 0.94 ± 0.04 | 0.46 ± 0.17 | 1.00 ± 0.00 | 1.00 ± 0.00 |
| $Na^+$-absent | 0.94 ± 0.03 | 0.9 ± 0.04 | 0.98 ± 0.01 | 1.00 ± 0.00 |

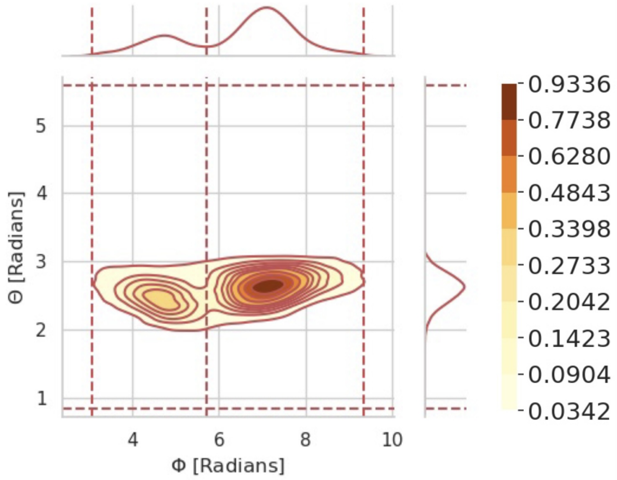


**Figure S 1.** Example of water molecule polarization distribution obtained from PENSA.

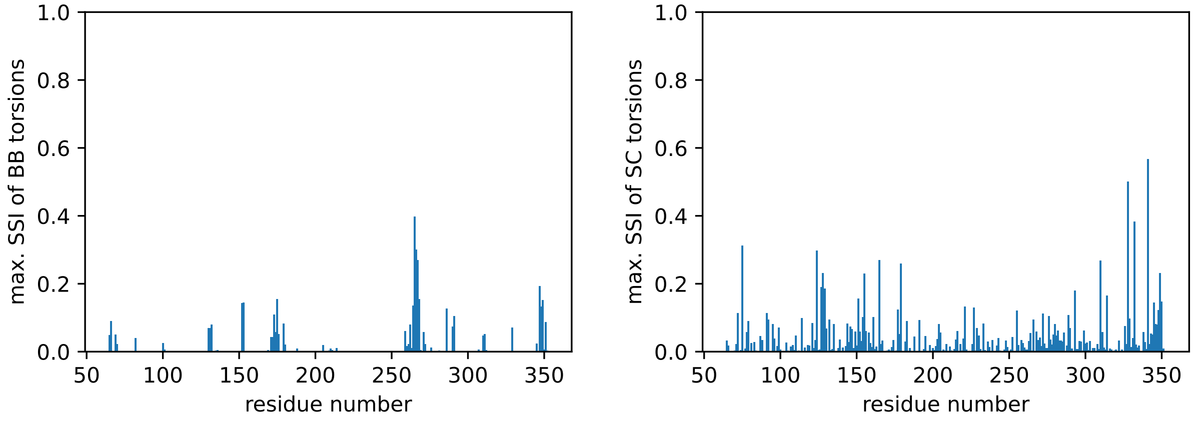


**Figure S 2.** State-Specific Information values for all receptor residue backbone (BB) and sidechain (SC) torsions, with each residue specified according to the sequence position.

| | | W1 | W2 | W3 | W4 |
|---|---|---|---|---|---|
| SSI [bits] | Polarisation | 0.02 | 0.00 | 0.41 | 0.64 |
| | Occupation | 0.00 | 0.17 | 0.00 | 0.00 |

**Table S 2.** State-Specific Information values for each intracavity water site, separated into the contributions from polarisation and occupation.
